# Deletion of the protein tyrosine phosphatase PTPN22 for adoptive T cell therapy facilitates CTL effector function but promotes T cell exhaustion

**DOI:** 10.1101/2023.06.24.546361

**Authors:** Alexandra R. Teagle, Patricia Castro-Sanchez, Rebecca J. Brownlie, Nicola Logan, Simran S. Kapoor, David Wright, Robert J. Salmond, Rose Zamoyska

## Abstract

**Background:** Adoptive cell therapy (ACT) is a promising strategy for treating cancer, yet it faces several challenges such as lack of long term protection due to T cell exhaustion induced by chronic TCR stimulation in the tumor microenvironment. One benefit of ACT, however, is that it allows for cellular manipulations, such as deletion of the phosphotyrosine phosphatase non-receptor type 22 (PTPN22), which improves CD8^+^ T cell anti-tumor efficacy in ACT. We tested whether *Ptpn22^KO^* cytolytic T cells (CTL) were also more effective than *Ptpn22^WT^* CTL in controlling tumors in scenarios that favor T cell exhaustion.

**Methods:** Tumor control by *Ptpn22^WT^* and *Ptpn22^KO^* CTL was assessed following adoptive transfer of low numbers of CTL to mice with subcutaneously implanted MC38 tumors. Tumor infiltrating lymphocytes were isolated for analysis of effector functions. An *in vitro* assay was established to compare CTL function in response to acute and chronic re-stimulation with antigen-pulsed tumor cells. The expression of effector and exhaustion-associated proteins by *Ptpn22^WT^* and *Ptpn22^KO^* T cells was followed over time *in vitro* and *in vivo* using the ID8 tumor model. Finally, the effect of PD-1 and TIM-3 blockade on *Ptpn22^KO^* CTL tumor control was assessed using monoclonal antibodies and CRISPR/Cas9-mediated knockout.

**Results:** Despite having improved effector function at the time of transfer, *Ptpn22^KO^* CTL became more exhausted than *Ptpn22^WT^* CTL, characterized by more rapid loss of effector functions, and earlier and higher expression of inhibitory receptors (IRs), particularly the terminal exhaustion marker TIM-3. TIM-3 expression, under the control of the transcription factor NFIL3, was induced by IL-2 signaling which was enhanced in *Ptpn22^KO^* cells. Anti-tumor responses of *Ptpn22^KO^* CTL were improved following PD-1 blockade *in vivo*, yet knockout or antibody-mediated blockade of TIM-3 did not improve but further impaired tumor control, indicating TIM-3 signaling itself did not drive the diminished function seen in *Ptpn22^KO^* CTL.

**Conclusions:** This study questions whether TIM-3 plays a role as an IR and highlights that genetic manipulation of T cells for ACT needs to balance short term augmented effector function against the risk of T cell exhaustion in order to achieve longer term protection.

**What is already known on this topic:**

- T cell exhaustion in the tumor microenvironment is a major factor limiting the potential success of adoptive cell therapy (ACT) in the treatment of solid tumors.
- Deletion of the phosphatase PTPN22 in CD8^+^ T cells improves their response to tumors, but it is not known whether this influences development of exhaustion.

**What this study adds:**

- Under conditions which promote exhaustion, CTL lacking PTPN22 exhaust more rapidly than WT cells, despite displaying enhanced effector function in their initial response to antigen.
- *Ptpn22^KO^* CTL express high levels of the inhibitory receptor TIM-3, but TIM-3 signaling does not directly contribute to *Ptpn22^KO^* CTL dysfunction.
- *Ptpn22^KO^* T cells are more responsive to IL-2 through JAK-STAT signaling, which induces TIM-3 expression via the transcription factor NFIL3.

**How this study might affect research, practice or policy:**

- Strategies aimed at augmenting T cell effector function for ACT should balance improved responses against an increased risk of T cell exhaustion.

## BACKGROUND

In recent decades, the field of cancer immunotherapy has expanded greatly, leading to improved outcomes for many patients. T cell-based immunotherapy in particular has yielded significant benefits, with great success delivered by immune checkpoint inhibitors, now used widely to treat numerous cancers(1). Despite these successes, not all patients benefit, with only a small proportion experiencing durable response. Multiple challenges preventing universal success are presented by the immunosuppressive tumor microenvironment (TME; reviewed in(2)). In recent years, it has been well established that a major obstacle to successful cancer immunotherapy is T cell exhaustion. This refers to a state of dysfunction in T cells in response to persistent stimulation with antigen in chronic viral infections and cancer, characterized by transcriptional and epigenetic changes leading to progressive loss of effector function, impaired persistence, and co-expression of multiple inhibitory receptors (IRs)(3).

Adoptive cell therapy (ACT), in which autologous peripheral blood or tumor infiltrating T lymphocytes (TILs) are manipulated *ex vivo* before expansion and infusion to the patient, provides an opportunity to surmount some of these barriers. Chimeric antigen receptor (CAR) expressing T cells are an example of ACT that has demonstrated remarkable results in patients with hematological malignancies refractory to conventional therapies(4). However, treatment of solid tumors with CAR T cells is yet to yield such promising results, in part because CARs recognize intact tumor cell surface proteins, while most solid tumor antigens are intracellularly derived peptides presented on the cell surface by MHC. Genetic engineering of autologous T cells to express αβ T cell receptors (TCRs) specific to such antigens is therefore an alternative approach for ACT in solid tumors. This approach also allows for additional modifications aimed at improving T cell effector function and longevity in tumors. It has been demonstrated previously that the autoimmunity-associated tyrosine phosphatase PTPN22 restrains T cell responses to weak affinity and/or self-antigens(5), and that systemic inhibition or deletion of PTPN22 improves responses to tumors(6,7). PTPN22 deletion restricted to T cells is sufficient to induce enhanced tumor control, as demonstrated by tumor rejection after adoptive transfer of *Ptpn22^KO^* naïve, effector or memory T cells into tumor bearing wild type host mice(8,9).

An important factor in developing a T cell product for optimal anti-cancer protection is the choice of T cell phenotype. Effector cytotoxic T lymphocytes (CTL) are straightforward to generate and expand to large numbers *in vitro* and have the greatest cytotoxicity, but their restricted longevity has raised questions about their ability to provide long-term tumor control. Prior investigation in *in vivo* tumor models has shown superior tumor control from adoptive transfer of *Ptpn22^KO^* CTL when administered in high numbers(8,9). However, systematic evaluation of the longer term fate of *Ptpn22^KO^* CTL in the face of persisting tumor challenge has not been carried out. Here, we use *in vitro* and *in vivo* models of chronic antigen exposure and show that despite *Ptpn22^KO^* CTL out-performing *Ptpn22*-sufficient (*Ptpn22^WT^*) CTL in their initial responses, CTL lacking PTPN22 more rapidly acquire an exhausted phenotype. In consequence, low numbers of *Ptpn22^KO^* CTL transferred to tumor-bearing mice were less effective than *Ptpn22^WT^* CTL at controlling established tumor growth and showed enhanced expression of inhibitory receptors, such as PD-1, and in particular TIM-3. Inhibition of PD-1 improved tumor control by *Ptpn22^KO^* CTL, supporting a previous report(7). In contrast, deletion or blockade of TIM-3 was detrimental to *Ptpn22^KO^* CTL function, suggesting TIM-3 does not function as a conventional inhibitory receptor in *Ptpn22^KO^* CTL and, therefore, is not a good target for reversing exhaustion in *Ptpn22^KO^* cells. Together, our findings illustrate that strategies aiming to optimize T cell effector function, such as PTPN22 deletion or inhibition, need to carefully consider selection of T cell phenotype for ACT in order to balance enhanced short-term effector function with susceptibility to exhaustion so as to optimize long-term tumor control.

## RESULTS

### PTPN22^KO^ CTL have enhanced effector function, but control tumors less effectively following adoptive transfer

To understand the impact of PTPN22 deletion in CTL for long term tumor control, we employed the Class I H-2K^b^-restricted OT-I TCR transgenic system, which allows TCR stimulus strength to be altered through use of cognate OVA peptides with varying affinities(10). CTL were generated from naïve OT-I *Rag1^KO^* T cells (hereafter referred to as OT-I cells), which were either *Ptpn22^WT^* or *Ptpn22^KO^*, by activating *in vitro* with the strong agonist peptide, SIINFEKL (N4), for 2 days prior to expansion in IL-2 for 4 days(5). In keeping with previous work(5,9), *Ptpn22^KO^* OT-I CTL produced more cytokine than *Ptpn22^WT^* OT-1 cells in response to 4 hours of re-stimulation with weak OVA peptide SIITFEKL (T4, Fig. 1a). In addition, *Ptpn22^KO^* OT-I CTL were ∼3-fold more cytotoxic than *Ptpn22^WT^* CTL against MC38 tumor cells expressing T4 (MC38-T4; Fig. 1b).

**Figure 1:**
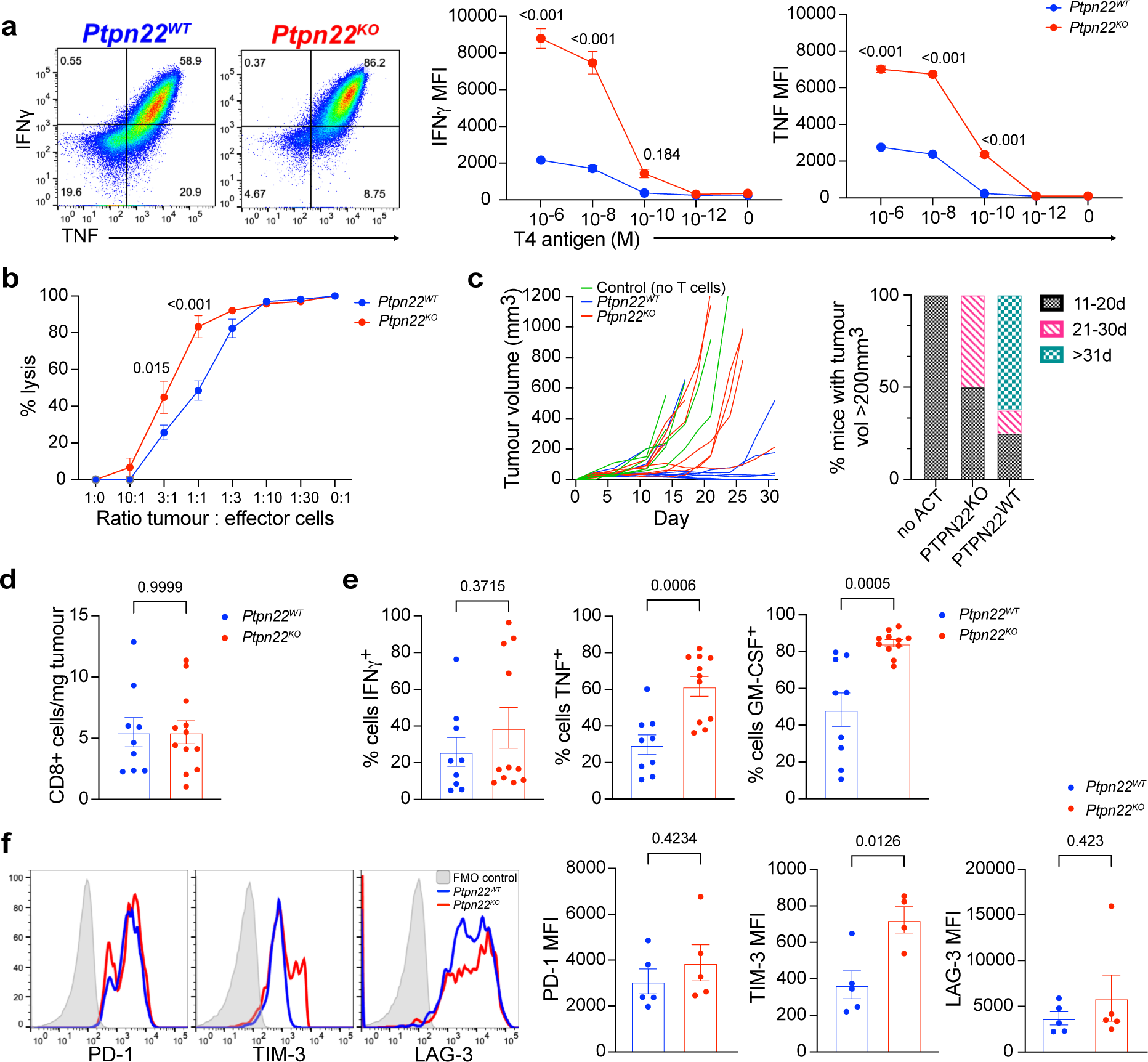
*Ptpn22^KO^* CTL are more effector-like, but control tumours less efficiently following adoptive transfer. a) Cytokine production from *Ptpn22^WT^* and *Ptpn22^KO^* CTL following re-stimulation for 4h with 10nM T4 (dot plots) or T4 at concentrations shown (graphs). Dot plots are gated on live, single cells. Numbers in dot plot quadrants are percentages. Data are representative of 4 independent experiments. b) Cytotoxicity of *Ptpn22^WT^* and *Ptpn22^KO^* CTL against MC38 tumour cells expressing T4 antigen, at ratios indicated. Data are representative of 4 independent experiments. c) MC38-T4 tumour growth (left) and time to tumours reaching 200mm^3^ volume in Rag1^KO^ hosts following ACT with *Ptpn22^WT^* and *Ptpn22^KO^* CTL, or no ACT. 0.5 x 10^6^ tumour cells were injected at day 0, followed by ACT with 1 x 10^6^ CTL at d4. n=5-8 per group in the experiment shown. Data are representative of 3 independent experiments (total n = 27 mice per group for *Ptpn22^WT^* and *Ptpn22^KO^* CTL; 10 per group for no ACT). d) Tumour infiltration by adoptively transferred *Ptpn22^WT^* and *Ptpn22^KO^* CTL. Mice were culled once tumours reached humane end points, and tumours dissociated to obtain single cell suspensions for flow cytometric analysis. Data are pooled from 2 independent experiments. n=9-12 per group. e) Cytokine production by *Ptpn22^WT^* and *Ptpn22^KO^* TIL. Mice were injected with Brefeldin A 4h before culling. Data are pooled from 2 independent experiments. Data were excluded from tumours with insufficient (<150) numbers of CD8+ TIL. n=9-12 per group. f) Inhibitory receptor expression on *Ptpn22^WT^* and *Ptpn22^KO^* TIL. Data are representative of 2 independent experiments. All bars on graphs represent mean ± SEM. p values as determined by 2-way ANOVA with Šidák correction for multiple comparisons (a, b), or Student’s t test (d, e, f). MFI; median fluorescence intensity.

Control of tumor growth by ACT is effectively a competition between tumor bulk and efficacy of the transferred T cells. Previous studies showed that transfer of large numbers (10^7^) of *Ptpn22^KO^* CTL provided better control of EL4 lymphoma and ID8 ovarian carcinoma than *Ptpn22^WT^* CTL(8,9). We asked whether this would also be the case if fewer cells were adoptively transferred, thus taking longer to gain control of the tumors, which might be more in keeping with a therapeutic scenario in which tumors are more established at the time of commencing treatment. MC38 tumor cells expressing the low affinity peptide, T4, were inoculated subcutaneously (s.c.) into *Rag1^KO^* mice, providing a model system in which tumor rejection would be mediated by the transferred CTL alone without contribution from the *Ptpn22^WT^* host T cell response. 10^6^ *Ptpn22^KO^* or *Ptpn22^WT^* CTL per recipient were injected intravenously once tumors were palpable, and tumor growth was followed. Unexpectedly, although better than no ACT, *Ptpn22^KO^* ACT controlled tumor growth less efficiently compared with mice receiving *Ptpn22^WT^* ACT (Fig. 1c). To understand the reduced efficacy of *Ptpn22^KO^* CTL in this situation, we analyzed tumors from the mice and found that both genotypes were equal in their ability to infiltrate and persist in tumors (Fig. 1d). Furthermore, assessment of cytokine production *in vivo* following Brefeldin A administration i.v. showed more *Ptpn22^KO^* than *Ptpn22^WT^* TILs stained positively for cytokines (Fig. 1e). However, expression of inhibitory receptors (IRs) such as PD-1 and LAG-3 was marginally higher on *Ptpn22^KO^* TILs, while expression of TIM-3 was significantly higher when compared to *Ptpn22^WT^* TILs (Fig. 1f), suggesting that heightened negative regulatory signals in *Ptpn22^KO^* TILs may play a role in their impaired control of tumors. Collectively, these results show that lack of PTPN22 in CTL enhances effector function in response to antigen, but if administered to tumor bearing hosts in numbers insufficient to rapidly control tumor growth, *Ptpn22^KO^* CTL are more prone to exhaustion and fail to give prolonged protection against tumors.

### PTPN22^KO^ CTL become more dysfunctional upon chronic TCR stimulation

To understand the reasons underlying the impaired tumor control by lower numbers of *Ptpn22^KO^* CTL, we modelled *in vitro* the chronic TCR stimulation experienced by TILs by repeatedly culturing CTL with antigen-pulsed tumor cells (Fig. 2a). As expected, d6 *Ptpn22^KO^* CTL were more cytotoxic and produced more cytokine in response to 4h re-stimulation with fresh T4-pulsed tumor cells (Fig. 2b). However, the functionality of both *Ptpn22^WT^* and *Ptpn22^KO^* CTL diminished upon repeated antigen exposure and by d9 of culture both genotypes lost the capacity to produce cytokines (Fig. 2b). Tumor target cell killing was still demonstrable up to d15 of culture by both *Ptpn22^KO^* and *Ptpn22^WT^* CTL, however the cytotoxic advantage exhibited by *Ptpn22^KO^* CTL at d6 (Fig 1b) was lost in favor of *Ptpn22^WT^* CTL at the later timepoint (Fig. 2c). While LAG-3 was highly expressed at similar levels by both cell genotypes, there was higher expression of IRs PD-1, TIGIT and most strikingly TIM-3 in *Ptpn22^KO^* compared to *Ptpn22^WT^* CTL at d6, even in the absence of antigen re-exposure (Fig. 2d, closed histograms); this was further boosted by 4h re-stimulation (Fig. 2d, open histograms). Over the time course of chronic re-stimulation PD-1 and TIGIT expression was consistently and significantly higher on *PTPN22^KO^* CTL, whilst TIM-3 expression was strikingly elevated on *Ptpn22^KO^* CTL throughout, yet its expression on *Ptpn22^WT^* cells remained low (Fig. 2e).

**Figure 2:**
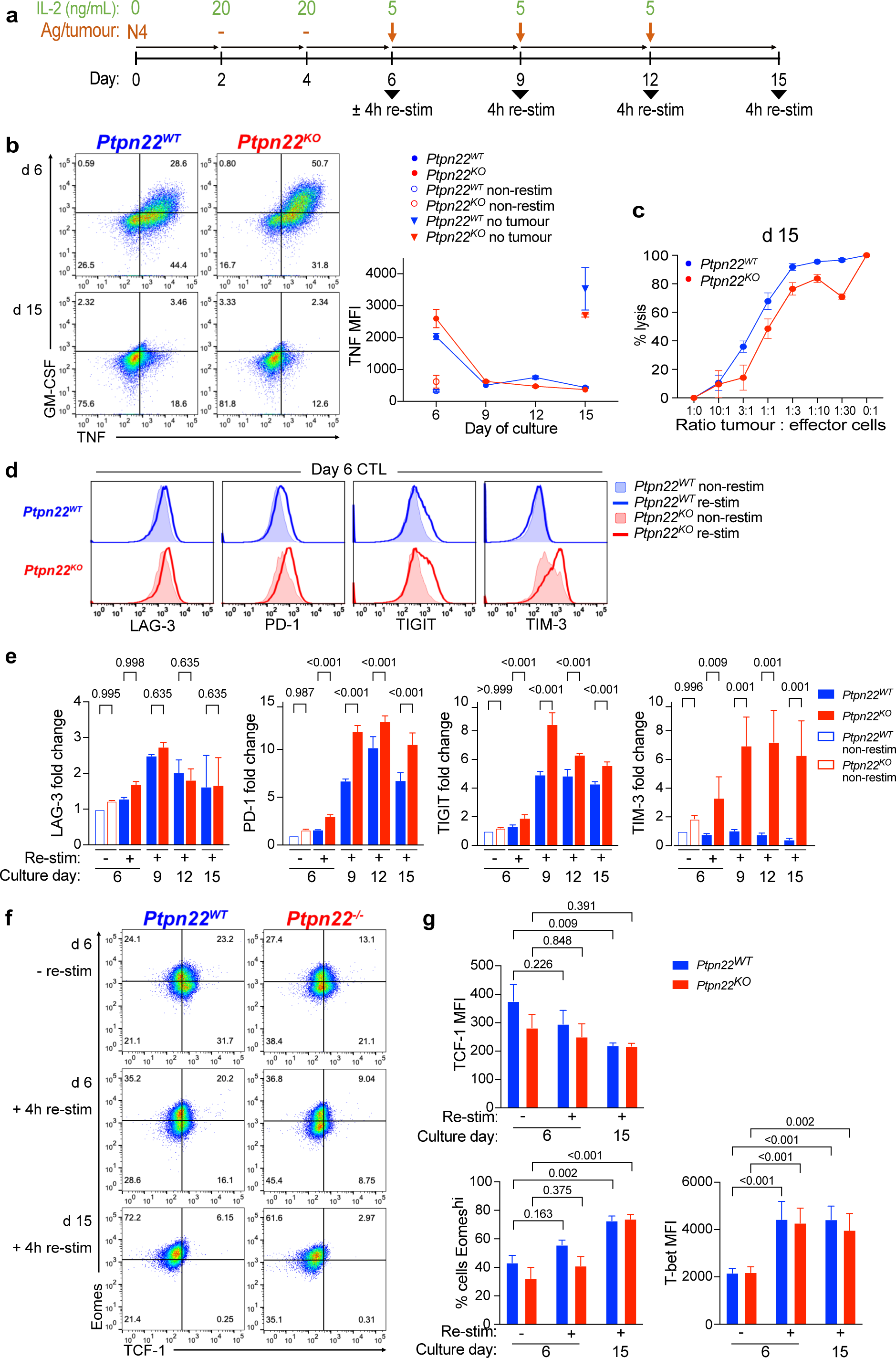
*Ptpn22^KO^* CTL become more dysfunctional upon chronic TCR stimulation. a) Schematic of experimental design. Naïve OT-I *Ptpn22^WT^* and *Ptpn22^KO^* T cells were activated with N4 (10nM) for 48 h, then cultured in IL-2 (20ng/mL) for 4d to expand them and induce differentiation to CTL. CTL on d6 were cultured with antigen-pulsed MC38 tumour cells, which were replenished every 3d. Cytokine production was assessed on cells either non-stimulated or re-stimulated for 4h with peptide as indicated. b) Cytokine production by *Ptpn22^WT^* and *Ptpn22^KO^* CTL following chronic re-stimulation with antigen. CTL were cultured for the indicated periods of time with MC38 tumour cells pulsed with N4 antigen. At each time point, CTL were re-stimulated with MC38 pulsed with T4 (100uM) for 4h and cytokine production in response was measured by intracellular staining for flow cytometry. Numbers in dot plot quadrants are proportions; gates are based on non-restimulated CTL. Graph shows TNF as representative cytokine. Data are representative of 3 independent experiments. c) Killing of luciferase expressing MC38-T4 cells by *Ptpn22^WT^* and *Ptpn22^KO^* CTL after chronic (d15) re-stimulation with antigen-bearing tumour cells. Data are representative of 3 independent experiments. d) Inhibitory receptor expression on *Ptpn22^WT^* and *Ptpn22^KO^* CTL, resting or re-stimulated for 4h with MC38 tumour cells pulsed with T4 (100uM). Data are representative of 3 independent experiments. e) Inhibitory receptor expression on *Ptpn22^WT^* and *Ptpn22^KO^* CTL after chronic re-stimulation with antigen. Data are pooled from two independent experiments. f) Eomes and TCF-1 expression by *Ptpn22^WT^* and *Ptpn22^KO^* over chronic Ag exposure. Representative dot plots of 2 independent experiments. g) TCF-1, Eomes and Tbet expression in *Ptpn22^WT^* and *Ptpn22^KO^* CTL at the indicated timepoints. Graphs show pooled data from 2 independent experiments. All bars on graphs represent mean ± SEM. p values as determined by 2-way ANOVA with Šidák correction for multiple comparisons (e, g). MFI; median fluorescence intensity.

Impaired function was accompanied by a change in abundance of certain transcription factors (TF), in particular T cell factor 1 (TCF-1; encoded by *Tcf7*) and Eomesodermin (Eomes), both of which are essential in T cell fate decisions, such as the generation of long-lived memory T cells(11–13) and these TF are characteristically changed in exhausted T cells(14,15). TCF-1 is essential for generation and maintenance of the CD8^+^ T cell memory response(11,13), as well as being associated with a stem-like population of exhausted cells in models of chronic infection and cancer(16–20). We found that expression of TCF-1 diminished with increasing tumor antigen exposure in keeping with reducing stemness and memory formation (Fig. 2f, g). Similarly Eomes, another transcription factor that is upregulated in exhausted T cells and which correlates with high IR expression(15) was increased following repeated Ag exposure (Fig. 2f, g). Expression of these transcription factors and of Tbet (Fig. 2g) was not significantly different between *Ptpn22^WT^* and *Ptpn22^KO^* cells at any timepoint, suggesting either a combined effect of several TFs rather than the action of one specifically, or a different TF was responsible for inducing the more dysfunctional and exhausted phenotype of cells lacking PTPN22.

### PTPN22^KO^ CTL become more dysfunctional in the tumor microenvironment

To understand how the response of *Ptpn22^KO^* CD8^+^ T cells developed *in vivo* we used the faster growing *p53^KO^* version of the ID8 ovarian carcinoma(21) transfected with N4 peptide (ID8-N4) which allows tumor responsive CD8^+^ T cells to be readily retrieved from the peritoneal exudate (PE). To eliminate any confounding effect of differing host environments we co-transferred a mix of naïve *Ptpn22^WT^* (CD45.1) and *Ptpn22^KO^*(CD45.1/2) OT-I T cells to C57BL/6 CD45.2 wild-type recipient mice with established tumors. Mice were culled at weekly intervals and CD8^+^ T cells in the PE and mesenteric lymph nodes (mLN) were analyzed (Fig. 3a). The presence of tumors was monitored in individual animals using *in vivo* bioluminescence imaging so that mice that rejected tumors could be excluded from the analysis, thus ensuring that only CD8^+^ T cells that had been continually exposed to tumor antigens were analyzed. Transferred T cell were readily detected in both PE and mLN at early time points (d4 and d12 post-transfer of T cells) but their numbers diminished with increasing time from transfer. Despite starting as 35% of the initial injection mix, *Ptpn22^KO^* T cell recovery was greater than that of *Ptpn22^WT^* T cells in peritoneal exudate at d4 and significantly more in both PE and mLN at d12 (Fig. 3b, c), indicating greater expansion in the initial response to antigen. However, by d19 the numbers of recovered *Ptpn22^WT^* and *Ptpn22^KO^* cells had equalized and by d26 *Ptpn22^KO^* cells were barely detectable, whilst low numbers of *Ptpn22^WT^* cells could still be identified in both mLN and PE. Notably, *Ptpn22^WT^* and *Ptpn22^KO^* T cells that were transferred into control mice without tumors were maintained and readily retrievable in the mLN at the later d26 timepoint (Fig. 3b, c). These findings show that T cells lacking PTPN22 initially undergo greater expansion in response to tumor antigens, however, when tumors are not cleared and thus antigen persists, *Ptpn22^KO^* cells that are chronically exposed to antigen *in vivo* fail to sustain proliferation and/or die more rapidly. This is specifically antigen dependent, since *Ptpn22^KO^* T cell survival was not impaired in tumor-free hosts.

**Figure 3:**
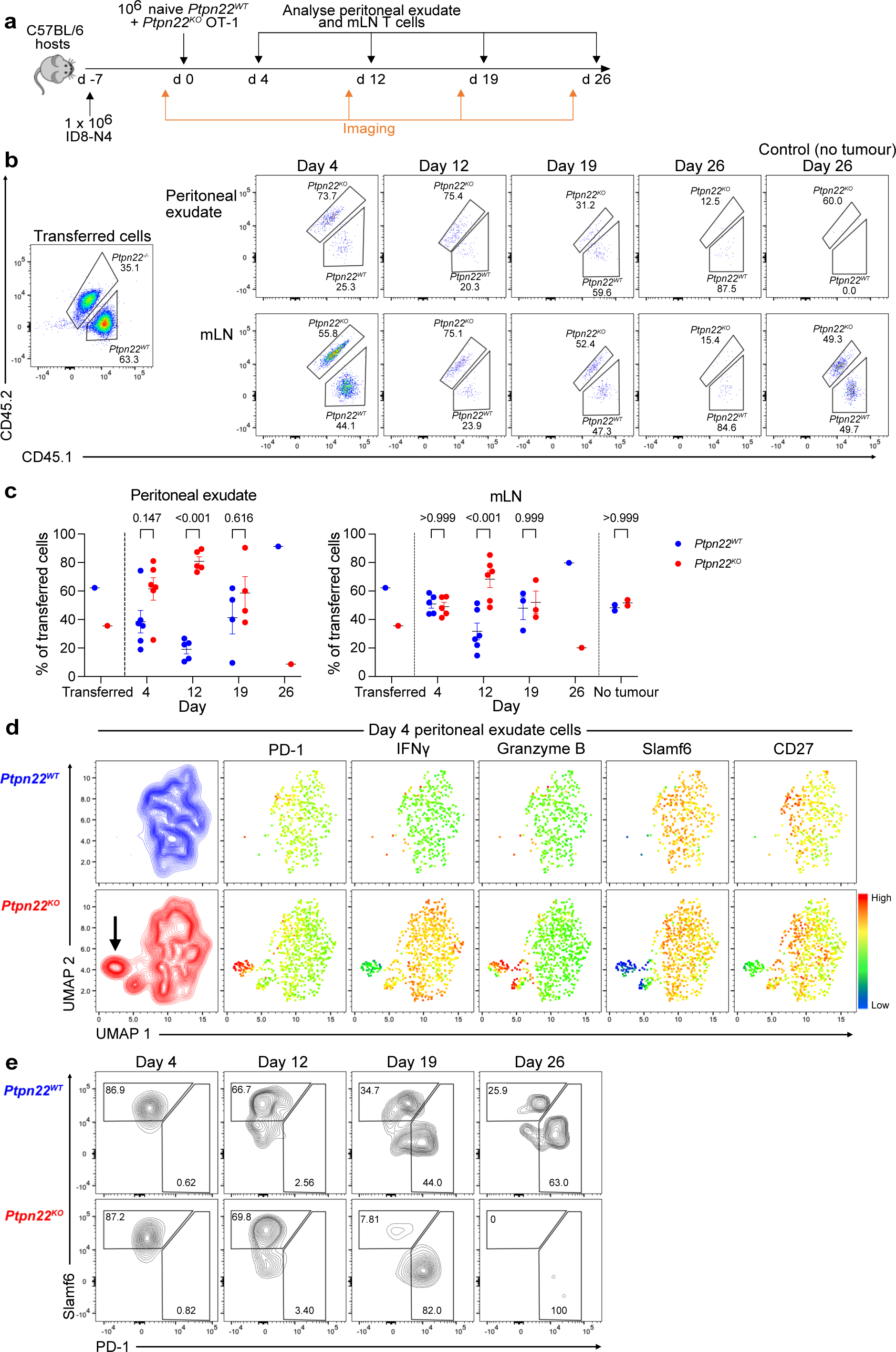
*Ptpn22^KO^* CTL become more dysfunctional in the tumour microenvironment. a) Experimental design. C57BL/6 hosts were injected i.p. with 1×10^6^ ID8-N4 tumour cells on day −7. IVIS imaging was carried out on day −1, prior to adoptive transfer of a mix of 1×10^6^ each of naïve *Ptpn22^WT^* and *Ptpn22^KO^* OT-I T cells. Tumour presence was confirmed with IVIS on the day before analysis of peritoneal exudate T cells at the indicated time points. Mice that spontaneously rejected tumours were excluded from analysis. n=6 mice at each time point; n=3 control mice (no tumour). b-c) Transferred *Ptpn22^WT^* and *Ptpn22^KO^* T cells in peritoneal exudate and mLN. b) Representative dot plots from each timepoint, as indicated. Gated on single, live, CD8^+^, CD45.1^+^ cells. Numbers are proportions of total donor cells. Control mice received T cells but no tumours. c) *Ptpn22^WT^* and *Ptpn22^KO^* cells in peritoneal exudate and mLN at indicated timepoints, as a proportion of total donor (CD45.1^+^) cells. Data are representative of 2 independent experiments. p values as determined by 2-way ANOVA with Šidák correction for multiple comparisons. d) UMAP embedding analysis of flow cytometry data showing donor *Ptpn22^WT^* and *Ptpn22^KO^* T cells in peritoneal exudate (gated on single, live, CD8^+^ CD45.1^+^ cells). Data from donor (CD45.1^+^) cells in all 6 mice at d4 were concatenated for UMAP analysis. Plots show expression of indicated proteins. e) Representative contour plots showing PD-1 and Slamf6 expression on *Ptpn22^WT^* and *Ptpn22^KO^* T cells isolated from peritoneal exudate of mice at the indicated time points (gated on single, live, CD8^+^ cells). Numbers in gates represent proportions.

In order to further characterize differences between *Ptpn22^WT^* and *Ptpn22^KO^* cells in tumor microenvironments, we performed uniform manifold approximation and projection (UMAP)(22) on flow cytometry data from *Ptpn22^WT^* and *Ptpn22^KO^* cells recovered from PE of mice with ID8 tumors at d4, when phenotypic changes were likely to be established. *Ptpn22^KO^* and *Ptpn22^WT^* cells occupied largely similar regions of phenotypic space but there were small distinct regions that were unique. In particular we identified a subpopulation of *Ptpn22^KO^* cells that separated from the bulk of the cells (Fig. 3d, indicated by arrow). This subpopulation expressed highly markers associated with terminal effector differentiation and exhaustion, such as PD-1 and Granzyme B. In addition, IFNγ was reduced suggesting early loss of effector function. In contrast the expression of markers associated with stemness or memory formation and longevity, including Slamf6 and CD27, was low. The subpopulation was absent in *Ptpn22^WT^* cells at day 4, suggesting that *Ptpn22^KO^* cells undergo terminal effector differentiation and exhaustion more readily than *Ptpn22^WT^* cells, potentially at the expense of memory cell differentiation.

We followed expression of Slamf6 and PD-1 on *Ptpn22^WT^* and *Ptpn22^KO^* cells from PE over the entire time-course of tumor exposure as indicators of stemness and terminal effector/exhaustion, respectively. These markers have been used to differentiate progenitor (Slamf6^hi^ PD-1^int^) and terminally exhausted (Slamf6^−^ PD-1^hi^) T cell populations in chronic viral infection and tumor models(14,23). Both *Ptpn22^WT^* and *Ptpn22^KO^* cells displayed a Slamf6^hi^PD-1^−^ phenotype at early timepoints consistent with a less differentiated state. However, Slamf6 expression was progressively lost and PD-1 acquired at later timepoints, in keeping with increasing terminal differentiation and development of exhaustion. Importantly, this occurred earlier in *Ptpn22^KO^* cells than in *Ptpn22^WT^* cells, with most *Ptpn22^KO^* cells identified as Slamf6^−^ PD-1^hi^ by d19 (Fig. 3e). These data are consistent with a model in which deletion of PTPN22 in T cells leads to acutely enhanced effector differentiation and function but this is at the expense of memory formation, and that in the setting of persisting antigen (such as in tumors) *Ptpn22^KO^* T cells ultimately exhaust more quickly than equivalent *Ptpn22^WT^* cells. This process is cell intrinsic and occurs even when both genotypes are exposed to the same environmental factors.

### Dysfunctional PTPN22^KO^ CTL can be rescued by PD-1 blockade

Given that IRs such as PD-1 were significantly elevated on *Ptpn22^KO^* CTL *in vitro* prior to adoptive transfer, we sought to test the hypothesis that this could be driving their impaired function. Previous studies have shown a synergistic effect of combining PTPN22 deletion or pharmacological inhibition with blockade of the PD-1/PD-L1 axis(6,7), but not the extent to which this treatment was due to a T cell-intrinsic effect excluding contributions from other hematopoietic PTPN22-expressing lineages. To determine the effect of PD-1 inhibition on *Ptpn22^KO^* CTL specifically, we combined our *Rag1^KO^* adoptive transfer model with PD-1 blocking mAb treatment (Fig. 4a).

**Figure 4:**
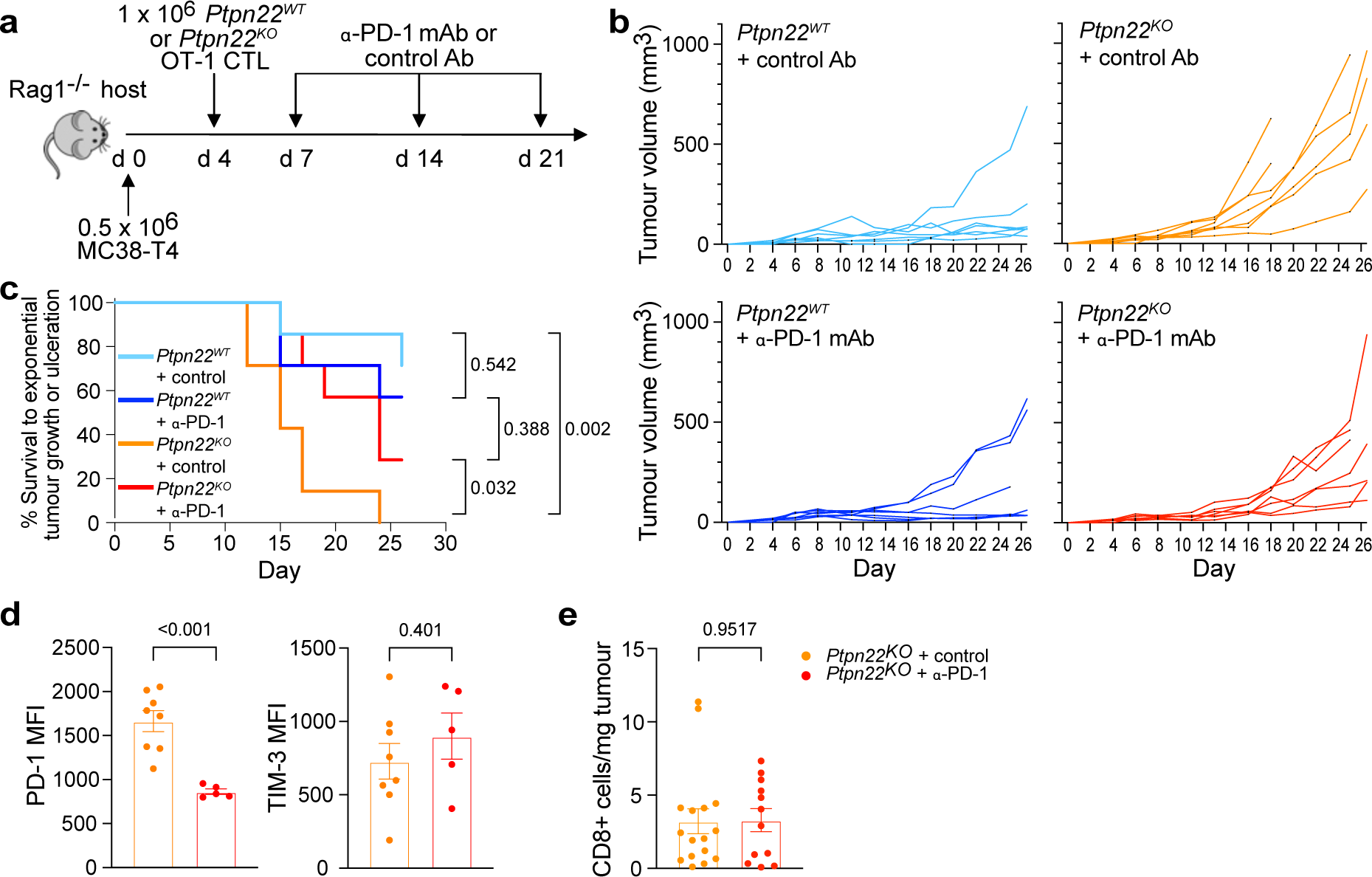
Dysfunctional *Ptpn22^KO^* CTL can be rescued by PD-1 blockade. a) Schematic of experiment. 0.5×10^6^ MC38-T4 tumour cells were injected at d0, followed by ACT with 1 x 10^6^ *Ptpn22^WT^* or *Ptpn22^KO^* CTL at d4, then i.p. injection of anti-PD-1 or isotype control Ab (200μg per mouse) on d7, d14, and d21. b) Tumour growth after ACT of *Ptpn22^WT^* or *Ptpn22^KO^* CTL ± anti-PD-1 mAb. Data are representative of 3 independent experiments. n=7 mice per group in the experiment shown (total n across all experiments=19-24). c) Survival to exponential tumour growth or ulceration. d) PD-1 and TIM-3 expression on *Ptpn22^KO^* TIL. Data are representative of 3 independent experiments. Tumours with fewer than 100 CD8^+^ TIL were excluded from analysis. n=6-9 per group in each experiment. e) Tumour infiltration by adoptively transferred *Ptpn22^KO^* CTL with or without anti-PD-1 mAb. Mice were culled once tumours reached humane end points, and tumours dissociated to obtain single cell suspensions for flow cytometric analysis. Data are pooled from 2 independent experiments. n=6-9 per group in each experiment. Bars on graphs represent mean ± SEM. p values as determined by pairwise survival analysis (b), or Student’s t test (d, e). MFI; median fluorescence intensity.

As before, adoptively transferred *Ptpn22^KO^* CTL provided less effective control of tumor growth, and this was significantly improved by anti-PD-1 treatment (Fig 4b, c). PD-1 blockade had no significant impact on tumor control by *Ptpn22^WT^* CTL, which were anyway effectively controlling tumor growth in these experiments. Further analysis of TILs from these mice indicated that the predominant effect of PD-1 mAb on *Ptpn22^KO^* TILs was to reduce surface PD-1 expression without significantly altering expression of other IRs such as TIM-3 (Fig. 4d) or infiltration and persistence in tumors (Fig. 4e). These data together provide further evidence that *Ptpn22^KO^* CTL become dysfunctional secondary to chronic antigen exposure in the tumor microenvironment, and that this dysfunction is reversible, albeit partly, by blockade of the PD-1 axis.

### TIM-3 inhibition worsens PTPN22^KO^ CTL tumor control

Alongside PD-1 expression, we noted that TIM-3 abundance was markedly increased on *Ptpn22^KO^* CTL (Fig. 1f, 2d, 2e). TIM-3 (encoded by *Havcr2*) is postulated to be an IR, although its precise role and signaling is less well defined than that of more classic IRs such as PD-1. The majority of evidence, particularly from human tumor data, points to an inhibitory role for TIM-3(24–28), and it is well documented that TIM-3 marks terminally exhausted cells in chronic viral infections and cancer(16,19,20,23). With this in mind and given the significant improvement by PD-1 blockade of *Ptpn22^KO^* CTL tumor control, we investigated the consequence of TIM-3 inhibition in *Ptpn22^KO^* cells. *Ptpn22^WT^* cells were not included in these analyses as they expressed very little TIM-3.

Electronic gating of FACS plots showed that *Ptpn22^KO^* CTL with highest TIM-3 expression had highest cytokine production *in vitro*, consistent with greater effector potential (Fig. 5a, b). We next sought to block TIM-3 signaling to investigate its role in *Ptpn22^KO^* CTL. We used CRISPR-Cas9 to knock out *Havcr2* in activated *Ptpn22^KO^* cells before differentiating them to CTL and performing functional assays *in vitro* (Fig. 5c). Two independent CRISPR guides were tested which gave similar KO efficiencies (Fig. 5d). In response to re-stimulation with antigen for 4h, we found that loss of TIM-3 did not significantly alter CTL function in terms of cytotoxicity (Fig. 5e) or cytokine production (Fig. 5f).

**Figure 5:**
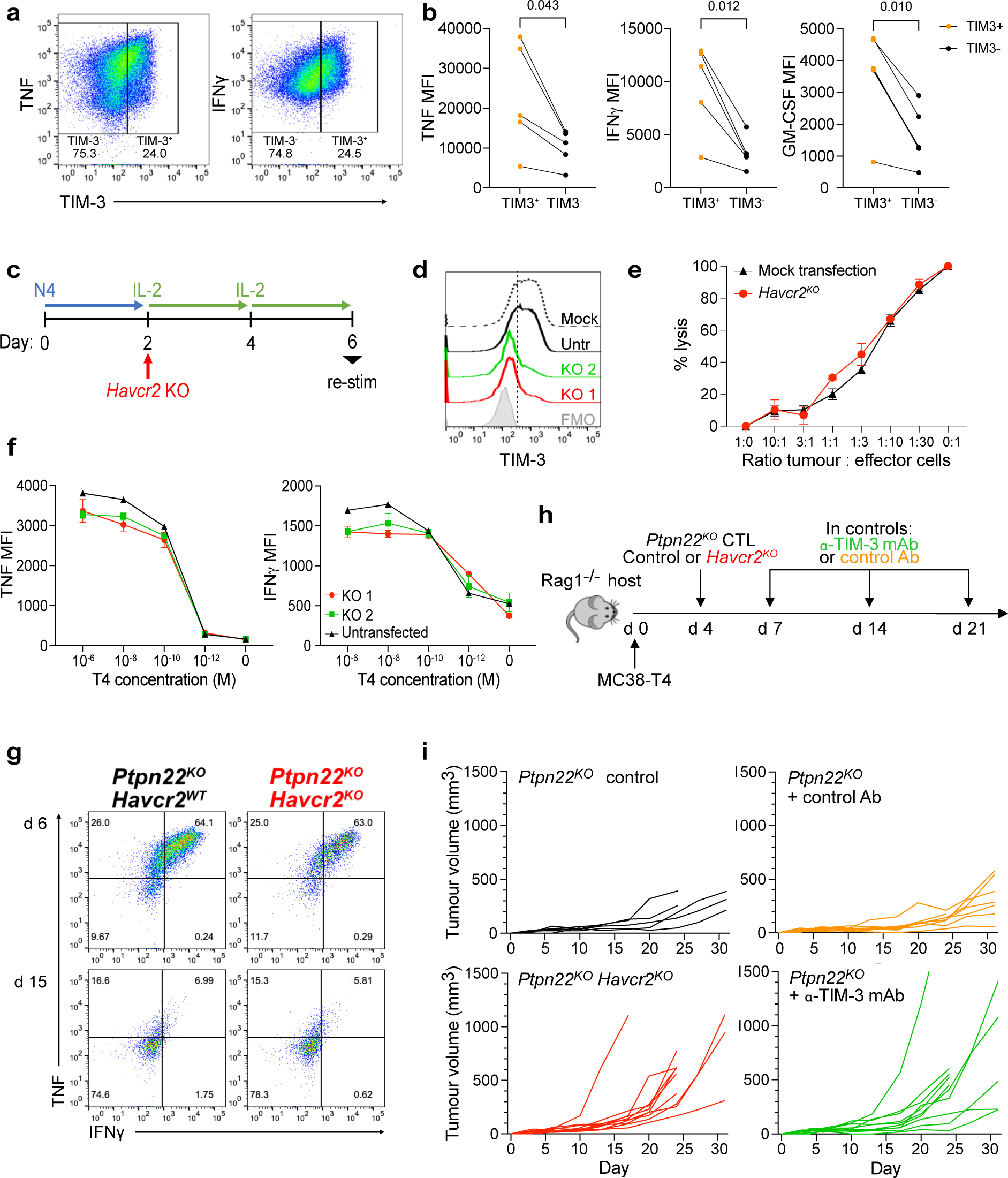
TIM-3 expression does not impair PTPN22^KO^ CTL function. a) TNF and IFNγ production by *Ptpn22^KO^* CTL following 4h re-stimulation with 10nM T4 antigen. Cells were gated on single live cells. Numbers in dot plot quadrants are proportions. Data are representative of 3 independent experiments. b) Cytokine production by *Ptpn22^KO^* CTL following 4h re-stimulation with 10nM T4 antigen. Data are pooled from 3 independent experiments. p values as determined by paired t test. c) Schematic of experimental design. Naïve OT-I *Ptpn22^KO^* T cells were activated with N4 (10nM) for 48h, then CRISPR-Cas9 was used to delete *Havcr2*, before cells were cultured in IL-2 (20ng/mL) for 4d to induce differentiation to CTL. d) Representative histograms showing TIM-3 expression on non-restimulated *Ptpn22^KO^* CTL on d6. Two independent guide RNA were tested (KO1 and KO2) in separate populations of cells. Untr; untransfected. e) Cytotoxicity of *Havcr2^KO^* KO or mock transfected *Ptpn22^KO^* CTL against MC38 tumour cells expressing T4 antigen, at ratios indicated. Data are representative of 2 independent experiments. Bars represent mean ± SEM. f) Cytokine production from *Ptpn22^KO^ Havcr2^KO^* and *Ptpn22^KO^* untransfected (*Havcr2^WT^*) CTL after 4h re-stimulation with T4 at indicated concentrations. Data are representative of 3 independent experiments. Bars represent mean ± SD. g) Representative dot plots showing cytokine production by *Ptpn22^KO^ Havcr2^+/+^* and *Ptpn22^KO^ Havcr2^KO^* CTL after 4h restimulation at d6 and chronic (d15) re-stimulation. Following *Havcr2* KO on d2 (Fig. 5c), cells were differentiated to CTL and repeatedly re-stimulated with antigen-bearing tumour cells as in Fig. 2a. Cells were gated on single, live, CD8^+^. Numbers in quadrants represent proportions. Data are representative of 2 independent experiments. h) Schematic of experiment. 0.5×10^6^ MC38-T4 tumour cells were injected at d0, followed by ACT with 1×10^6^ *Ptpn22^KO^* control or *Ptpn22^KO^ Havcr2^KO^* CTL at day 4. Groups of mice that had received control T cells were given anti-TIM-3 or isotype control Ab on d7, d14, and d21. i) Tumour growth in groups as in (h). n=5-10 mice per group in experiment shown. Data are representative of 2 independent experiments (total =10-20 per group). MFI; median fluorescence intensity.

We asked whether an impact of TIM-3 signaling might only be revealed over a more prolonged period or in response to chronic TCR stimulation, such as in the tumor microenvironment. First, we tested *Havcr2^KO^* CTL in chronic re-stimulation assays *in vitro* (as indicated in Fig. 2a). However, *Havcr2^KO^/Ptpn22^KO^* and *Havcr2^WT^*/*Ptpn22^KO^* CTL developed an equivalent loss of function following chronic antigen exposure (Fig. 5g). Second, to establish whether there was an influence of TIM-3 on *Ptpn22^KO^* CTL function in tumors, we adoptively transferred *Havcr2^WT^* or *Havcr2^KO^ Ptpn22^KO^* CTL into mice with MC38-T4 tumors. Additionally, for a group of mice that received control (*Havcr2^WT^*) cells, we administered TIM-3 blocking or isotype control monoclonal antibody (Fig. 5h). Strikingly, loss of TIM-3 either via knockout or mAb-mediated blockade impaired control of tumors by *Ptpn22^KO^* CTL (Fig. 5i), suggesting a positive benefit from the presence of TIM-3 on these dysfunctional cells in preserving some control against tumors. Together these data suggest that the high expression of TIM-3 on *Ptpn22^KO^* CTL is not a driver of their impaired function in response to chronic antigen exposure and, contrary to expectation, TIM-3 upregulation may be a compensatory mechanism to maintain a degree of function in the face of exhaustion in the tumor microenvironment.

### *Ptpn22^KO^* T cells have increased IL-2 signaling, which amplifies TIM-3 expression

Although TIM-3 blockade did not reverse *Ptpn22^KO^* CTL dysfunction, in view of the marked difference in its expression between *Ptpn22^WT^* and *Ptpn22^KO^* CTL, we sought instead to exploit TIM-3 as an indicator of the drivers leading to *Ptpn22^KO^* CTL dysfunction. Evaluation of IR expression at various timepoints during differentiation from naïve to effector CTL in *Ptpn22^KO^* cells revealed that TIM-3 is regulated differently to other “classic” IRs. PD-1 and TIGIT were induced by the initial TCR activation and boosted by subsequent TCR re-stimulations (Fig. 2a; days 0-2 and day 6 + re-stim), with their expression declining when cells were maintained in IL-2 alone (Fig. 2a; days 2-6; Fig. 6a). In contrast TIM-3 was not upregulated in response to TCR activation in naïve cells. Instead, TIM-3 cell surface expression was detected upon expansion of *Ptpn22^KO^* cells in IL-2 (Fig. 2a; days 2-6), with a subsequent boost in expression following TCR re-stimulation with antigen (Fig. 6a). TIM-3 expression in *Ptpn22^WT^* OT-1 cells remained low under these conditions.

**Figure 6:**
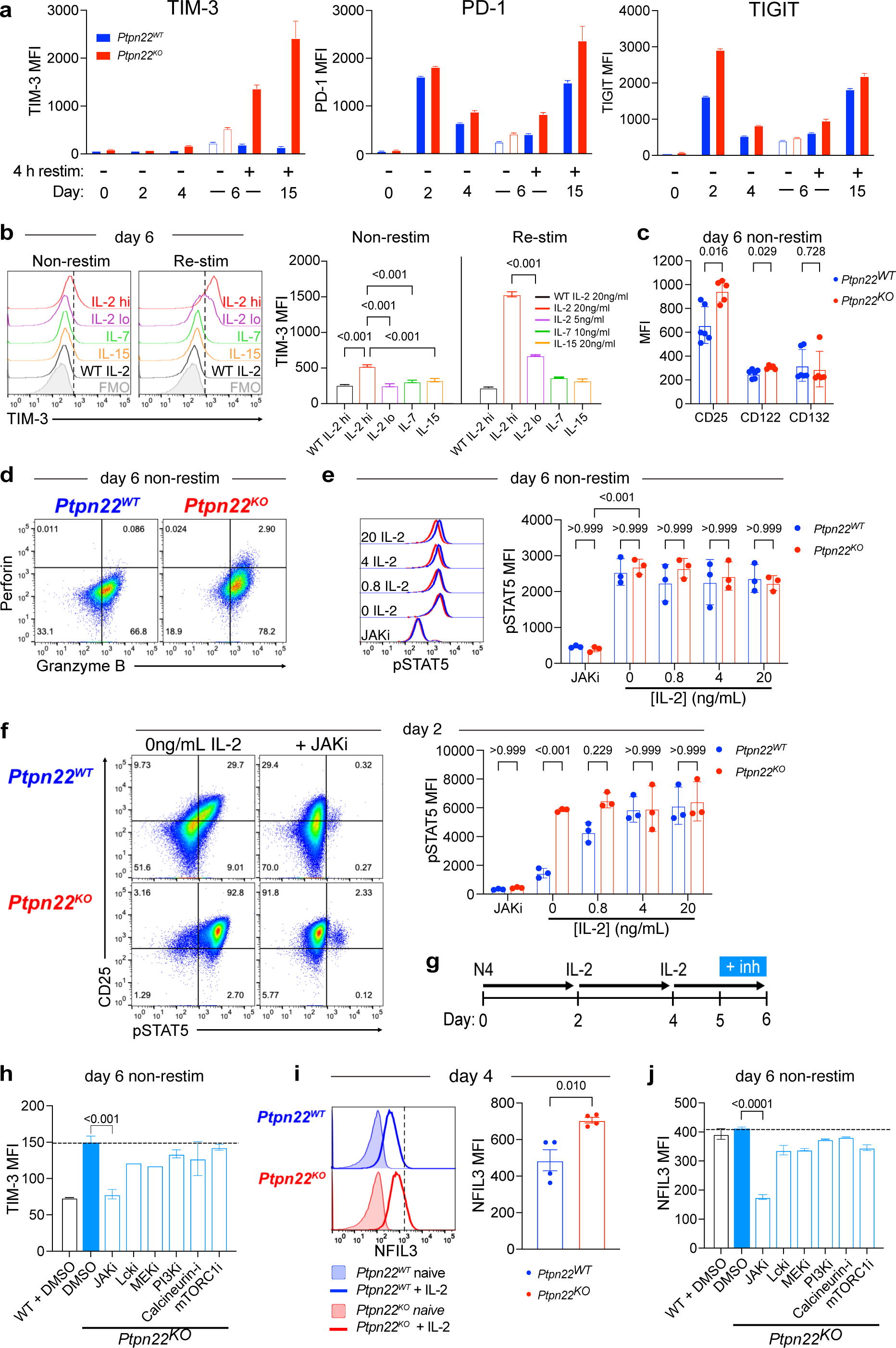
*Ptpn22^KO^* T cells have increased IL-2 signalling, which induces TIM-3 expression. a) OT-I *Ptpn22^WT^* or *Ptpn22^KO^* T cells were differentiated to CTL and then repeatedly stimulated with antigen-pulsed tumour cells as in Fig. 2a. Inhibitory receptor expression was measured by flow cytometry at the indicated time points. Data are representative of at least 3 independent experiments. b) Naïve OT-I *Ptpn22^WT^* or *Ptpn22^KO^* T cells were activated with N4 (10nM) for 48 hours, then cultured in the indicated cytokine for a further 4d: IL-2 20ng/mL (IL-2 hi); IL-2 5ng/mL (IL-2 lo); IL-7 10ng/mL; IL-15 20ng/mL. On d6, expression of TIM-3 was analysed by flow cytometry on resting (non-restim) cells, or after 4h re-stimulation with 10nM N4 peptide (re-stim). Data are representative of 3 independent experiments. c) IL-2 receptor chain expression by non-restimulated *Ptpn22^WT^* or *Ptpn22^KO^* CTL on d6. Data are combined from 3 independent experiments. d) Production of perforin and granzyme B by non-restimulated *Ptpn22^WT^* or *Ptpn22^KO^* CTL on d6, following 4d culture in IL-2 containing media (20ng/mL). Representative dot plots from one of 3 independent experiments. Gates are based on fluorescence minus one (FMO) controls. Numbers in quadrants are proportions. e) STAT5 phosphorylation in *Ptpn22^WT^* or *Ptpn22^KO^* CTL on d6. Cells were removed from culture on d6, washed and rested for 30 min in fresh media alone (0ng/ml IL-2), or with added JAK inhibitor tofacitinib, or with added IL-2 at the concentrations indicated. Data shown are from 3 biological replicates and are representative of 2 independent experiments. f) STAT5 phosphorylation in *Ptpn22^WT^* or *Ptpn22^KO^* T cells on d2 (after activation with N4 Ag). Cells were removed from culture, washed and rested for 30min in fresh media alone (0ng/mL IL-2), or with added JAK inhibitor tofacitinib, or with added IL-2 at the concentrations indicated. Data shown are from 3 biological replicates and are representative of 3 independent experiments. g) Schematic of experimental design. *Ptpn22^WT^* or *Ptpn22^KO^* T cells were differentiated to CTL as described previously. Inhibitors were added for the final 24 h of culture in IL-2. h) TIM-3 expression on *Ptpn22^KO^* CTL (d6) after culture in the presence of inhibitors stated during the final 24h of culture in IL-2. Dotted line indicates TIM-3 MFI in cells treated with vehicle control only. Data are representative of 3 independent experiments. Comparisons without p values shown did not reach significance. i) NFIL3 expression in *Ptpn22^WT^* or *Ptpn22^KO^* T cells on d4, after activation and then 48h culture in IL-2 (20ng/ml), shown relative to NFIL3 in naïve cells in histogram. Data in graph are pooled from 2 independent experiments. j) NFIL3 expression in *Ptpn22^KO^* CTL after culture in the presence of inhibitors stated during the final 24h of culture in IL-2. Dotted line indicates NFIL3 MFI in cells treated with vehicle control only. Data are representative of 2 independent experiments. Comparisons without p values shown did not reach significance. Bars on graphs represent mean ± SD (a, c, e, f, h, j), or mean ± SEM (b, i). p values as determined by one-way ANOVA with Dunnett’s multiple comparisons test (b, h, j), or Student’s t test (c, i), or two-way ANOVA with Šidák correction for multiple comparisons (e, f). MFI; median fluorescence intensity.

Induction of TIM-3 on *Ptpn22^KO^* cells was specific to culture in IL-2, in a dose dependent manner. These data concur with a previous report that TIM-3 is one of the proteins most reduced in abundance when CTL are deprived of IL-2(29). Other cytokines such as IL-7 and IL-15 which also signal through the common gamma chain (γ_c_ cytokines) were unable to induce TIM-3 expression, even when the cells were subsequently re-stimulated with antigen (Fig. 6b). Indeed, *Ptpn22^KO^* CTL had increased expression of CD25 (IL-2Rα) as well as CD122 (IL-2Rβ), components of the IL-2R complex that are normally limiting for responsiveness to IL-2(30), but not CD132 (IL-2Rγ) (Fig. 6c), suggesting increased responsiveness to IL-2. In support of this, we also found higher expression of IL-2 targets(29) such as perforin and granzyme B in *Ptpn22^KO^* CTL (Fig. 6d).

To explore whether *Ptpn22^KO^* T cells were more responsive to IL-2, we measured phosphorylation of the downstream signaling molecule STAT5. STAT5 phosphorylation was equivalent in *Ptpn22^WT^* and *Ptpn22^KO^* CTL at day 6 (Fig. 6e). However, we reasoned that by this timepoint cells had been exposed to high concentrations of exogenous IL-2 for a prolonged period which may have plateaued STAT5 phosphorylation in both genotypes. In contrast, at day 2 of culture STAT5 phosphorylation was significantly greater in *Ptpn22^KO^* cells that had been activated with antigen for 48h compared to *Ptpn22^WT^* counterparts (Fig. 6f), suggesting increased IL-2 signaling in *Ptpn22^KO^* cells and indicating that this is initiated early in differentiation by enhanced responsiveness to antigen in the absence of PTPN22. To confirm this was an IL-2 specific response, *Ptpn22^KO^* and *Ptpn22^WT^* T cells were cultured with peptide for 48h, then washed into fresh media to remove any secreted cytokines and rested in cytokine-free media for 30 minutes prior to incubation with a titration of IL-2 for 30 mins. Interestingly, activated *Ptpn22^KO^* cells rested in fresh media without supplemental IL-2 retained significantly higher levels of phospho-STAT5 than *Ptpn22^WT^* cells, which was inhibitable in both genotypes by JAK inhibitor. Addition of IL-2 at concentrations at or above 4ng/ml for 30 minutes equalized the pSTAT5 signal between *Ptpn22^KO^* and *Ptpn22^WT^* T cells suggesting that *Ptpn22^KO^* cells intrinsically produce and utilize IL-2 to a greater extent than *Ptpn22^WT^* cells.

To validate these findings, we used specific small molecule inhibitors to block pathways downstream from the IL-2 receptor (Fig. 6g). Only the JAK inhibitor, tofacitinib, reduced TIM-3 expression to a level similar to that of WT CTL (Fig. 6h), confirming a critical role for JAK-STAT signaling in regulating IL-2-induced TIM-3 expression in *Ptpn22^KO^* T cells. The fact that IL-2 was the only γ_c_ cytokine to induce TIM-3 expression suggested that IL-2 high affinity receptor binding induces JAK-STAT5 signaling to an extent that surpasses a threshold not attained by IL-15 or IL-7, and our data indicate that in *Ptpn22^KO^* cells this threshold is more readily attained with increased STAT5 phosphorylation in response to initial antigen stimulation. Previous studies have shown that the transcription factor NFIL3 induces TIM-3 expression downstream from the IL-2 receptor(29,31), and we confirmed that NFIL3 expression was greater in *Ptpn22^KO^* than in *Ptpn22^WT^* T cells following culture in IL-2 (Fig. 6i), and that IL-2-induced NFIL3 upregulation is sensitive to JAK inhibition with tofacitinib (Fig. 6j).

These data show that enhanced responsiveness to antigen in *Ptpn22^KO^* T cells increases IL-2 production and expression of receptor components CD25 and CD122, which leads to increased STAT5 phosphorylation, setting up a positive feedback loop. This ultimately induces TIM-3 expression via JAK-STAT signaling and the transcription factor NFIL3, and likely contributes to their accelerated development of dysfunction in the context of chronic antigen stimulation, since IL-2 signaling promotes terminal effector differentiation(30,32,33) as well as exhaustion(34,35).

## DISCUSSION

CD8^+^ T cells have great potential as adoptive cell therapeutics in cancer, but improvements are needed to optimize their utility. The diversity of phenotypes of responding CD8^+^ T cells is of key importance for successful ACT, and must comprise a blend of short-lived effector cells to rapidly gain control of tumor mass, as well as long lived memory cells for more durable protection. *Ptpn22^KO^* T cells were previously shown to control tumor growth more efficiently(8,9) and, once tumors were cleared, formed memory populations that effectively limited tumor re-inoculation(9). Therefore, in the present study it was unexpected to find that transferred *Ptpn22^KO^* CTL were less able than *Ptpn22^WT^* CTL to control tumor growth. Despite exaggerated effector responses to brief antigen re-encounter, we show that chronic re-stimulation by tumor antigens led to a decline in *Ptpn22^KO^* CTL function. A key difference here was that a ten-fold lower inoculum of CTL was administered to tumor bearing mice compared to our previous studies, such that the resulting incomplete tumor clearance allowed us to follow the development of an exhaustion phenotype(36) associated with up-regulation of inhibitory receptors which correlated with impaired anti-tumor immunity(37). Our use of *Rag1^KO^* hosts could have impacted tumor control by transferred CTL, since CD4^+^ T cell help – which is absent in *Rag1^KO^* mice – is essential to sustain CTL activity; however, we found that the accelerated acquisition of an exhausted phenotype in *Ptpn22^KO^* T cells was replicated across different tumor models including after co-transfer of *Ptpn22^WT^* and *Ptpn22^KO^* T cells to the same WT hosts.

In terms of inhibitory receptors, most striking was the excessively high expression of TIM-3 by *Ptpn22^KO^* T cells under conditions of prolonged Ag and cytokine exposure. This finding provided an opportunity to explore the role and regulation of this postulated inhibitory receptor. Despite initially being suggested to play a role in T cells in regulating autoimmunity(38), TIM-3 lacks any classical inhibitory signaling motifs in its cytoplasmic domain and consequently opinion has been somewhat divided as to its role in T cells. Over-expression of TIM-3 in Jurkat cells led to augmented T cell activation due to enhanced TCR signaling(39), while in acute LCMV infection TIM-3 was found to promote short-lived effector T cell differentiation but impair memory precursor T cell differentiation(40). Despite this, the majority of evidence in recent years points to TIM-3 having (at least predominantly) inhibitory effects(38,41,42). In mouse models of cancer TIM-3 overexpression increased tumor progression(43), whereas blockade synergized with PD-1 blockade to improve tumor inhibition(44). Strong evidence for an inhibitory role also comes from human data, as severity and prognosis of many solid and hematological malignancies correlates negatively with TIM-3 expression(25,45–47). Moreover, TIM-3 positivity marks out the most dysfunctional cells in CD8^+^PD-1^+^ populations(44,48), and co-blockade of PD-1 and TIM-3 has been more effective than PD-1 blockade alone in improving function of T cells from patients with metastatic melanoma(48,49). In contrast, we found that loss of TIM-3 through either antibody-mediated blockade or deletion of the receptor further impaired *Ptpn22^KO^* CTL control of tumors. Despite the deficiency of self-renewal capacity or polyfunctionality in terminally exhausted T cells, of which TIM-3 is characteristic(16,19,20,23), it is this population that retains cytotoxicity and is responsible for tumor control(23). Therefore, it is possible that removing TIM-3 or blocking its ligand interactions in TIM-3^hi^ exhausted *Ptpn22^KO^* TILs depletes those remaining cells with limited but enduring cytotoxic potential, thereby further impairing their anti-tumor response. It is notable that Cubas *et al* similarly identified a greater proportion of PD-1^+^LAG-3^+^TIM-3^+^ CD8^+^ TILs in tumors from *Ptpn22^KO^* mice, and that these cells expressed greater levels of granzyme B, suggesting some enduring heightened cytotoxicity(6). TIM-3 has been shown to interact with both Lck(50) and Fyn(39) when it is bound or unbound, respectively, by ligand; since Lck and Fyn are substrates of PTPN22(51,52) there may be direct interaction between TIM-3 and PTPN22-mediated signaling. Further studies will be useful in elucidating any such relationships. Importantly, TIM-3 is found on other immune cells, including regulatory T cells(47), myeloid cells(53), natural killer (NK) cells(54), and mast cells(55), and thus previously demonstrated favorable responses to systemic TIM-3 blockade may be dependent on multiple interacting cell types. Indeed, several recent studies have established a critical role for dendritic cells in response to anti-TIM3 mAbs(56–58). Finally, the aforementioned reports of enhanced T cell function secondary to TIM-3 signaling(39,40) may suggest that its role is context-dependent. Interestingly, recently reported phase 1a/b clinical trials of anti-TIM3 mAbs in patients with various advanced solid tumors showed no or only minimal clinical benefit when the drugs were used as monotherapy(59,60), suggesting a more subtle and nuanced role for TIM-3 compared to other inhibitor receptors.

TIM-3 expression on *Ptpn22^KO^* T cells was upregulated by IL-2 but not other γ_c_ cytokines, indicating that only IL-2 surpasses a necessary threshold of JAK signaling, and that *Ptpn22^KO^* cells reach this threshold more readily than *Ptpn22^WT^* cells. The availability and abundance of IL-2 and whether it is produced and consumed in an autocrine manner or acquired extracellularly impacts memory versus effector differentiation(61), and IL-2 signaling has additionally been implicated in driving T cell exhaustion(34). Initial antigen stimulation causes *Ptpn22^KO^* T cells to make more IL-2 than *Ptpn22^WT^* counterparts, which is particularly apparent in response to weak antigens(8). We found that *Ptpn22^KO^* T cells subsequently have elevated phosphorylation of STAT5 and maintain higher expression of CD25 and CD122 (Fig. 6c), and as such are able to initiate and potentially sustain heightened responsiveness to IL-2 through JAK-STAT signaling. Such increased IL-2-JAK1/3-STAT5 signaling may in turn increase their propensity to exhaustion, since STAT5 was recently shown to mediate exhaustion, including upregulating TIM-3 expression via direct binding to the *Havcr2* locus(35). Given that IL-2 is the cytokine which is most efficient at expanding T cells and producing the very large numbers of CD8^+^ T cells required for ACT protocols, any manipulation, such as genetic deletion of PTPN22, that may improve ACT function also needs to be considered as a potential driver of enhanced terminal effector differentiation and increased susceptibility to exhaustion. This may also be relevant when considering a recent report that murine CAR T cells lacking PTPN22 were not superior to WT CAR T cells in clearance of various solid tumors(62). Importantly, we have previously shown that *Ptpn22^KO^* T cells can become functional memory cells with exposure to appropriate cytokines(9), however in conditions favoring effector cell differentiation, PTPN22-deficient cells are exaggerated in their response to antigen and IL-2, which drives them harder towards short-lived effector and exhausted fates.

Multiple phosphatases have been shown to be potential targets to improve anti-tumor responses by ACT(63) so it is important to understand what tips the balance between efficacy and exhaustion when a phosphatase is absent. We consider that deletion of PTPN22 remains a viable strategy for improving cancer immunotherapies such as adoptive cell therapy, however careful consideration needs to be given to the differentiation state of the targeted T cells, in order to balance short term augmented effector function with longer term protection. Ultimately, in the “arms race” between cancer cells and the immune system, adopting an approach which boosts multiple T cell phenotypes is likely to be preferable.

## METHODS

### Patient and Public Involvement

No patients were involved in this research study.

### Mice

Mice expressing the OT-1 TCR transgene (C57BL/6-Tg(TcraTcrb)1100Mjb/J) backcrossed to the Rag-1KO (B6.129S7-*Rag1^tm1Mom^*/J) background and containing congenic alleles for CD45.2 or CD45.1 were bred onto the *Ptpn22^KO^* background. OT-1hom *Rag1^KO^*, *Ptpn22^KO^* OT-1hom *Rag1^KO^, Rag1^KO^*, and C57BL/6 mice were maintained under specific pathogen-free conditions at Bioresearch and Veterinary Services facilities at the University of Edinburgh. Mice were age and sex matched for all experiments. For *in vivo* tumor experiments, recipient mice were additionally assigned to groups based on tumor size to allow equal mean tumor volume in each group at the point of adoptive T cell transfer.

### Cell lines

MC38 colon adenocarcinoma cells were obtained from Doreen Cantrell (University of Dundee). T4 (SIITFEKL) ova-variant peptide constructs (a kind gift from Dietmar Zehn) and firefly luciferase constructs (a kind gift from Hans Stauss) were each introduced by retroviral transduction (MC38-T4-luc). ID8 ovarian carcinoma cells expressing N4 (SIINFEKL) and firefly luciferase were obtained as previously described(9) and CRISPR/Cas9 was used to delete p53 (ID8-N4-fluc-p53^−/−-^) to enhance tumorigenicity. Cells were maintained in IMDM supplemented with 10% FCS, L-glutamine, 100 U penicillin and 100 mg/mL streptomycin.

### Tumor models

To obtain tumor cell suspensions, adherent cells were dissociated from culture flasks using trypsin-EDTA (Gibco^TM^), counted and resuspended in sterile PBS (Sigma-Aldrich). 0.5 x 10^6^ MC38 tumor cells in 20μL sterile PBS were injected s.c. into the right dorsal flank of 6-12 week old *Rag1^KO^* mice after shaving, or 1 x 10^6^ ID8 tumor cells in 100μL sterile PBS were injected i.p. to C57BL/6 mice. For MC38 experiments, tumors were measured with digital calipers on day 4, and mice were divided into groups based on an equal mean tumor volume in each group. Sample size was based on a difference in mean tumor size between groups of 20%, a standard deviation of 1.4, 80% power, and a type I error rate of 0.5%. Mice from different experimental groups were mixed in cages together to minimize potential confounders. For ID8 experiments, presence of tumors was confirmed with non-invasive in vivo bioluminescence imaging (IVIS) and Living Image Software (Perkin Elmer). Mice were injected i.v. with *Ptpn22^WT^* or *Ptpn22^KO^* T cells, as described in individual figure legends. Control mice received no T cells but were injected with PBS. For experiments blocking inhibitory receptors, 200 μg anti-mouse PD-1 mAb (clone RMP1-14; InVivoMab, Bio X Cell) or 250 μg anti-TIM-3 mAb (clone RMT3-23; InVivoMab, Bio X Cell) were injected i.p. on days 7, 14 and 21 of tumor growth. For MC38 tumor models, tumors were measured with calipers every 2-3 days, and volume was calculated using the formula *V = 0.5(L x W^2^).* Mice were removed from the experiment when tumor maximum diameter reached 15mm or tumors ulcerated. Tumors were resected before mechanical dissociation and TILs isolation using a Ficoll-Paque gradient. For ID8 tumor models, mice were culled in groups at the indicated timepoints in Figure 3 and peritoneal exudate was obtained by peritoneal wash with ice cold 1% BSA in PBS. For tumor growth experiments, tumors from mice with fewer than 100 TIL isolated were excluded from analysis of TIL. For experiments in fig. 3, mice that spontaneously rejected tumors were excluded, as determined *a priori*. Researchers were not blinded to the group allocations during the experiments or analysis.

### In vitro T cell culture and differentiation

For generation of effector CTLs, naïve T cells were isolated from OT-1 lymph nodes and stimulated with 10nM N4 peptide (Cambridge Peptides) for 2 days in IMDM supplemented with 10% FCS, L-glutamine, 100 U penicillin, 100 mg/mL streptomycin, and 50μM β-mercaptoethanol. At day 2, cells were washed and then expanded and differentiated in complete IMDM (as above) containing 20 ng/mL recombinant human IL-2 (Peprotech) for a further 4 days, with media and IL-2 being refreshed after 48 hours. For experiments using different cytokines (Figure 6), cells were cultured from day 2 as above, or instead in IL-2 (5ng/mL), IL-7 (10ng/mL) or IL-15 (20ng/mL).

### In vitro cytotoxicity assay

Target MC38-T4-luc cells were seeded in 96 well plates and allowed to settle and adhere overnight. T cells were added to wells in triplicate at ratios indicated in figures, and cells were incubated together at 37°C for 4 hours. Media was then removed and plates were washed gently in PBS to remove T cells and debris. Remaining tumor cells were lysed using passive lysis buffer (Promega) and bioluminescence activity of each well was measured using D-luciferin (Luciferase Assay System, Promega) and Varioskan™ microplate reader (Thermo Scientific). Percentage tumor lysis was calculated from bioluminescence of surviving tumor cells relative to control wells containing no effectors or no targets (corresponding to 0% lysis and 100% lysis, respectively). Where tumor cell lysis was >100% or <0%, this was normalized to 100% or 0%, respectively.

### In vitro chronic re-stimulation assay

CTL were generated *in vitro* as described above. From day 6, CTL were added to culture vessels containing irradiated MC38 tumor cells that had been pulsed with T4 or N4 antigen (100μM) for 1-2 hours and then washed. CTL were added to tumor cells at a tumor to effector ratio of 1:3. Every 3 days CTL were resuspended and removed from any remaining tumor cells, washed, counted, and added onto fresh irradiated tumor cells that had been pulsed with Ag as before. At days 6, 9, 12, and 15 of culture, a sample of CTL was taken to perform functional assays by re-stimulating with fresh pulsed MC38 cells in 96-well plates. Day 6 re-stimulation represents acute re-stimulation only, without chronic antigen exposure.

### Flow cytometry

The following anti-mouse antibodies were used. From Biolegend: CD8b-APC/Cy7 (clone YTS156.7.7) or CD8b-BV510 (clone YTS156.7.7), CD8a-BV421 (clone 53-6.7), IFNγ-PE/Cy7 (clone XMG1.2), TNF-PerCP/Cy5.5 (clone MP6-XT22), TIM-3-APC (clone B8.2C12), LAG-3-PE/Cy7 (clone C9B7W), TIGIT-BV421 (clone 1G9), CD45.1-BV605 (clone A20), CD45.2-BV750 (clone 104), Granzyme B-Pacific Blue (clone GB11), CD27-AF700 (clone LG.3A10), CD25-PerCP (clone PC61), Perforin-PE (clone S16009B), CD71-FITC (clone RI7217), CD98-PE/Cy7 (clone RL388). From Thermo Fisher Scientific: GM-CSF-PE (clone MP1-22E9), PD-1-FITC (clone RMP1-30), Eomes-PerCP-eFluor710 (clone Dan11mag), NFIL3-PE (clone S2M-E19). From BD Biosciences: PD-1-BV650 (clone J43), TCF-1-PE (clone S33 966), CD122-FITC (clone 5H4), CD132-PE (clone 4G3), pSTAT5-AF647 (clone 47), CD8a-PE (clone 53-6.7). From Miltenyi: Slamf6-FITC (clone 13G3). Live/dead-Aqua or live/dead-NIR dyes were used (Life Technologies). For *ex vivo* analysis of cytokines in TILs, mice were injected i.v. with Brefeldin A (Cambridge Bioscience) 4 hours prior to culling. For analysis of cytokines, granzyme B and perforin *in vitro*, cells were re-stimulated for 4 h with N4 or T4 peptides at the concentrations indicated in relevant figures, in the presence of Brefeldin A. Cells were labelled with live/dead and surface stains prior to fixation and permeabilization with fixation buffer (Biolegend) and intracellular staining permeabilization wash buffer (Biolegend). Alternatively, FoxP3 fixation/permeabilization buffers (eBioscience) were used for staining for perforin and for transcription factors. Samples were acquired using a MACS Quant analyzer 10 (Miltenyi), or Aurora spectral flow cytometer (Cytek) (Figure 3), and data were analyzed using FlowJo software (Treestar). Uniform manifold approximation and projection (UMAP) analysis was performed using the FlowJo plugin. For UMAP analysis (Fig. 3), cell populations were separated based on the following parameters: TIGIT (in BV421), Granzyme B (in Pacific Blue), CD44 (in BV570), PD-1 (in BV650), CD62L (in BV711), CD103 (in BV786), Slamf6 (in FITC), perforin (in PE), CD127 (in PE-dazzle 594), TNF (in PerCP-Cy5.5), 4-1BB (in PerCP-eF710), IFNγ (in PE-Cy7), KLRG1 (in PE-Fire810), TIM-3 (in APC) and CD27 (in AF700).

### CRISPR/Cas9 knock out of Havcr2

CRISPR/Cas9 technology was used to delete *Havcr2*, as previously reported(64), using the Neon™ Transfection system (Thermo Fisher Scientific) according to manufacturer instructions. Guide RNAs (gRNA) were purchased from Integrated DNA Technologies. TrueCut™ Cas9 (Thermo Fisher Scientific) was used. Five *Havcr2* targeting gRNA were trialed and the two giving the greatest transfection efficiency, as measured at protein level by flow cytometry and DNA disruption confirmed with PCR, were taken forward for experiments.

### Inhibitors

For experiments in figures 6g, h, and i, cells were treated with 1μM Tofacitinib (Stratech), 10μM PP2 (Sigma-Aldrich), 10μM U0126 (Promega), 10μM IC87114 (Sigma-Aldrich), 10μM Cyclosporin A (Sigma-Aldrich), or 200nM Rapamycin (Sigma-Aldrich). Control cells were treated with DMSO. Relevant inhibitors were added to culture media during the final 24 h of culture prior to analysis of cells. For experiments in figures 6e-f, cells were treated with 2μM tofacitinib.

### Statistics

Statistical analyses were performed using GraphPad Prism 9.5. Statistical analyses used are described in the relevant figure legends. *P* < 0.05 was considered significant. Error bars represent S.E.M. unless otherwise stated in figure legends.

## DECLARATIONS

### Ethics approval

This study was approved by the Ethical Review Body at the School of Biological Sciences, University of Edinburgh. All animal experiments were approved by the University of Edinburgh Bioresearch and Veterinary Services Ethical Review body and the United Kingdom Home office under project license P38881828 to RZ.

### Patient consent for publication

Not applicable.

### Availability of data and material

Data are available upon reasonable request.

### Authors’ contributions

ART designed and performed experiments, analyzed data, and wrote the manuscript. PCS and RJB designed and performed experiments, analyzed data, and contributed to development of the project. NL performed experiments in *in vivo* models. SK performed experiments and analyzed data. DW performed transduction of the MC38 cell line and optimized CRISPR-Cas9 experiments. RJS contributed to development of the project and writing of the manuscript. RZ contributed to conception and development of the project and experimental design, and writing of the manuscript. All authors read and approved the final version of the manuscript.

## Acknowledgments

We are grateful to D Cantrell for the provision of cells lines, and H Stauss and D Zehn for the provision of firefly luciferase constructs and ova-variant peptide constructs, respectively.

## LIST OF ABBREVIATIONS

ACT: Adoptive cell therapy
CAR: Chimeric antigen receptor
CTL: Cytotoxic T lymphocyte
i.p.: Intra-peritoneal
IR: Inhibitory receptor
i.v.: Intra-venous
mLN: mesenteric lymph node
PE: Peritoneal exudate
PTPN22: Protein tyrosine phosphatase non-receptor 22
s.c.: Sub-cutaneous
TCF-1: T cell factor 1
TCR: T cell receptor
TF: Transcription factor
TIL: Tumor infiltrating lymphocyte
TME: Tumor microenvironment
UMAP: Uniform manifold approximation and projection

